# Peripheral blood mononuclear cells preferentially activate 11-oxygenated androgens

**DOI:** 10.1101/2020.09.08.288316

**Authors:** Lina Schiffer, Alicia Bossey, Angela E Taylor, Ildem Akerman, Dagmar Scheel-Toellner, Karl-Heinz Storbeck, Wiebke Arlt

**Affiliations:** Institute of Metabolism and Systems Research, University of Birmingham, Birmingham, UK; Institute of Inflammation and Ageing, University of Birmingham, Birmingham, UK; Department of Biochemistry, Stellenbosch University, Stellenbosch, South Africa; National Institute for Health Research (NIHR) Birmingham Biomedical Research Centre, University Hospitals Birmingham NHS Foundation Trust and University of Birmingham, Birmingham, UK

**Author notes:** Corresponding author and contact for reprint requests: Professor Wiebke Arlt, Institute of Metabolism and Systems Research (IMSR), University of Birmingham, B15 2TT, Birmingham, UK.

**Keywords:** AKR1C3, 11-oxygenated androgens, androgen metabolism, liquid chromatography-tandem mass spectrometry, PBMCs, natural killer cells

## Abstract

**Context:** Androgens are important modulators of immune cell function impacting proliferation, differentiation and cytokine production. The local generation of active androgens from circulating androgen precursors is an important mediator of androgen action in peripheral androgen target cells or tissue.

**Objective:** To characterize the activation of classic and 11-oxygenated androgens in human peripheral blood mononuclear cells (PBMCs).

**Methods:** PBMCs were isolated from healthy male donors and incubated ex vivo with precursors and active androgens of the classic and 11-oxygenated androgen pathways. Steroids were quantified by liquid chromatography-tandem mass spectrometry. The expression of genes encoding steroid-metabolizing enzymes was assessed by quantitative PCR.

**Results:** PBMCs generated 8-fold higher amounts of the active 11-oxygenated androgen 11-ketotestosterone than the classic androgen testosterone from their respective precursors. We identified the enzyme AKR1C3 as the major reductive 17β-hydroxysteroid dehydrogenase in PBMCs responsible for both conversions and found that within the PBMC compartment natural killer cells are the major site of AKRC13 expression and activity. Steroid 5α-reductase type 1 catalyzed the 5α-reduction of classic but not 11-oxygenated androgens in PBMCs. Lag time prior to the separation of cellular components from whole blood increased 11KT serum concentrations in a time-dependent fashion, with significant increases detected from two hours after blood collection.

**Conclusions:** 11-oxygenated androgens are the preferred substrates for androgen activation by AKR1C3 in PBMCs, primarily conveyed by natural killer cell AKR1C3 activity, yielding 11KT the major active androgen in PBMCs. Androgen metabolism by PBMCs can affect the measurement results of serum 11-ketotestosterone concentrations, if samples are not separated in a timely fashion.

## Introduction

A critical step in androgen activation is the conversion of the androgen precursor androstenedione (A4) to the active androgen testosterone (T) (1-3), which occurs in the testes catalyzed by 17β-hydroxysteroid dehydrogenase type 3. However, in the majority of peripheral target tissues of androgen action, the intracrine activation of A4 to T is catalyzed by the enzyme AKR1C3 (aldo-keto reductase 1C3, also termed 17β-hydroxysteroid dehydrogenase type 5) (1-3), thereby representing a key regulator of local androgen exposure. Peripheral blood mononuclear cells (PBMCs) can activate A4 to T (4, 5) catalyzed by AKR1C3 (6). Androgens impact on a multitude of immune cell functions, including proliferation, differentiation, apoptosis, cytokine and immunoglobulin productions (7, 8) and also link immune and metabolic regulation (9).

Recent in vitro work has shown that AKR1C3 has an approximately 8-fold higher catalytic efficiency for the generation of the 11-oxygenated androgen 11-ketotestosterone (11KT) from its precursor 11-ketoandrostenedione (11KA4) than for the generation of T from A4 (10). 11KT activates the androgen receptor with potency and efficacy similar to T (11-13) and is the predominant circulating androgen in androgen excess conditions including congenital adrenal hyperplasia, polycystic ovary syndrome and premature adrenarche (12, 14, 15). 11KT is derived from the abundant adrenal precursor steroid 11β-hydroxyandrostenedione (11OHA4) (13, 16), which is converted to 11KA4 by peripheral tissue expressing 11β-hydroxysteroid dehydrogenase type 2 (17, 18) prior to activation to 11KT by AKR1C3 (10, 16).

Of note, a number of studies have shown gradual increases in testosterone measured in serum, if the blood sample is left unseparated from cellular components for an increased period of time, using either immunoassays (19-21) or ultra-high performance liquid chromatography-tandem mass spectrometry (LC-MS/MS) for steroid quantification (22, 23). One of those studies additionally investigated the effect of time before separation on 11KT concentrations measured in serum and observed a much more pronounced increase in 11KT than T over time (23), which could have a differential impact on the measurement of circulating 11-oxygenated androgen concentrations.

In this study, we explore this further and undertake a detailed characterization of androgen activation via both classic and 11-oxygenated androgen biosynthesis pathways in the human peripheral blood cell compartment and subpopulations, utilizing PBMCs isolated from healthy volunteers.

## Material and Methods

### Blood collection

The collection of blood for PBMC isolation and serum steroid analysis underwent ethical approval by the Science, Technology, Engineering and Mathematics Ethical Review Committee of the University of Birmingham (ERN_17-0494, ERN_17-0494A, and ERN_14-0446) and all participants gave informed, written consent prior to study participation. Exclusion criteria for participation were any acute or chronic disease affecting steroid biosynthesis or metabolism and intake of any medication known to interfere with steroid biosynthesis or metabolism. All blood samples were collected between 09:00h and 11:00h; 50 mL of blood were collected into K_2_ EDTA Vacutainers™ (purple top) for the immediate isolation of PBMCs.

To assess the effect of the time period between blood collection and separation of the cellular components by centrifugation on measured serum steroid concentrations, blood was collected into several SST™ Vacutainers™ (gold top). The tubes were left unseparated at room temperature in an air-conditioned room and were separated by centrifugation (2000*g*, 10 min) after defined time periods (0, 1, 2, 4, 6 and 24 hours). The separated serum was stored at -80 °C.

### Ex vivo steroid metabolism assays in peripheral blood mononuclear cells

PBMCs were isolated from blood by density gradient centrifugation using Ficoll Paque Plus (GE Healthcare) following the manufacturer’s instructions. Isolated PBMCs were washed in RPMI-1640 (Sigma-Aldrich) supplemented with 100 U/mL penicillin and 0.1 mg/mL streptomycin, counted and assessed for viability by trypan blue exclusion. A minimum of 3×10^6^ cells was frozen in 400 μL TRI reagent^®^ (Sigma Aldrich) and stored at -80 °C for RNA analysis at a later date. For steroid metabolism assays, 3×10^6^ cells were incubated with 100 nM of the respective steroid in a final volume of 500 μL RPMI-1640 supplemented with penicillin and streptomycin. Steroids were added from stock solutions in methanol and the final methanol concentration in the incubations was 0.00304% (v/v). Technical replicates were prepared, if sufficient cell numbers were available. During the incubation, the samples were constantly gently rotated at 37 °C for 24 hours. For each experiment, incubations of cells with methanol only and cell-free incubations for all steroids were prepared as controls as well as dilutions of each steroid at 100 nM in medium that were frozen at -20 °C immediately after preparation. At the end of the incubation period samples centrifuged for 2 minutes at 10,000 rpm. The supernatant was stored at -20 °C for steroid analysis. The cell pellet was washed in PBS, suspended in 50 μL lysis buffer (50 mM Tris pH 8.0, 150 mM NaCl, 0.1% SDS, 0.5% sodium deoxycholate, 1% Trition X-100, 0.1 mM DTE, 0.1 mM PMSF, 0.1 mM EDTA) and stored at -80 °C for protein quantification.

### Natural killer cell ex vivo steroid metabolism assays

PBMCs were prepared as described above and NK cells were subsequently enriched using the MACS human NK cell isolation kit (Miltenyi Biotec) as per the manufacturer’s instructions. Cells were checked for viability, counted and steroid conversion assays were set up as described above.

### Steroid analysis by tandem mass spectrometry

The following steroids were quantified by LC-MS/MS in the medium supernatant of the PBMC incubations and serum samples as previously described (15): 11β-hydroxy-5α-androstanedione (11OH-5α-dione, 5α-androstane-11β-ol-3,17-dione), 11β-hydroxyandrostenedione (11OHA4, 4-androstene-11β-ol-3,17-dione), 11β-hydroxytestosterone (11OHT, 4-androstene-11β,17β-diol-3-one), 11-ketoandrosterone (11KA4, 4-androstene-3,11,17-trione), 11-ketotestosterone (11KT, 4-androstene-17β-diol-3,11-dione), 5α-dihydrotestosterone (DHT, 5α-androstane-17β-ol-3-one), 5α-androstanediol (Adiol, 5α-androstane-3α,17β-diol), 5α-androstanedione (5α-dione, 5α-androstane-3,17-dione), androstenedione (A4, 4-androstene-3,17-dione), androsterone (An, 5α-androstane-3α-ol-17-one), dehydroepiandrosterone (DHEA, 5-androstene-3β-ol-17-one), testosterone (T, 4-androstene-17β-ol-3-one). In addition, 11-keto-5α-dihydrotestosterone (11KDHT, 5α-androstane-17β-ol-3,11-dione) and 11-keto-5α-androstanedione (11K-5α-dione, 5α-androstane-3,11,17-dione) were analyzed by ultra-high performance supercritical fluid chromatography-tandem mass spectrometry (UHPSFC-MS/MS) as previously described (24). For ex vivo cell incubations, steroid concentrations or their ratios were normalized to the total protein content of the incubations as determined in the supernatant after cell lysis using the DC Protein Assay (Bio-Rad).

### RNA extraction quantitative PCR

Samples stored in TRI reagent^®^ were defrosted and the RNA in the aqueous phase of the phenol-chloroform extraction was purified using the RNeasy Mini Kit (Qiagen). RNA concentrations were determined from the absorbance of the sample at 260 nm using a Nanodrop spectrophotometer and reverse transcription was performed using Applied Biosystems™ TaqMan™ Reverse Transcription Reagents following the manufacturer’s protocol. Quantitative PCR (qPCR) was performed on an ABI 7900HT sequence detection system (Perkin Elmer, Applied Biosystems) using TaqMan™ Gene Expression Assays (FAM-labelled) and the SensiFASTTM Probe Hi-ROX kit (Bioline). ΔCt was calculated as Ct [Target]-Geometric mean (Ct [HPRT1], Ct [GAPDH]). Gene expression in arbitrary units (A.U.) was calculated as 1000*2^-ΔCt. For targets not reproducibly detected in duplicate reactions, relative gene expression is shown as 0.

### Analysis of published RNAseq data from PBMC subpopulations

We accessed dice-database.org (25) to investigate *AKR1C3* expression determined by RNAseq in FACS-sorted PBMC subpopulations. The database contains expression data from 54 male and 37 female healthy donors (age range 18-61 years). *AKR1C3* expression data in log2 transformed transcripts per million (TPM) were downloaded and plotted in GraphPad Prism 8.

### Statistical analysis

Changes in serum steroid concentrations were analyzed in GraphPad Prism 8 using ANOVA followed by a Dunnett multiple comparison test to compare each timepoint against the sample processed immediately after collection (0h). Statistical analysis of differences in steroid concentrations in the supernatants from the PBMC ex vivo incubations was performed by Wilcoxon matched-pairs signed rank test. Statistical analysis of RNAseq data was performed by one-way ANOVA followed by Tukey’s multiple comparisons test or Mann-Whitney test as appropriate.

## Results

### Androgen activation in PBMCs is primarily catalyzed by AKR1C3 and SRD5A1

Using qPCR, we identified AKR1C3 as the major reductive, activating 17β-hydroxysteroid dehydrogenase isoform in PBMCs, while HSD17B3, generally considered a testes-specific isoform, was expressed at very low levels only (**Fig. 1A**). While we could not detect expression of steroid 5α-reductase type 2 (SRD5A2) and 17β-hydroxysteroid dehydrogenase type 2 (HSD17B2) in any of the samples from 14 donors using our qPCR assay, we detected consistently high expression of steroid 5α-reductase type 1 (SRD5A1) in samples from all donors (n=14) and detected 17β-hydroxysteroid dehydrogenase type 4 (HSD17B4) expression in samples from 6 of 14 donors. This indicates that SRD5A1 is responsible for the 5α-reduction we observe in our PBMC incubations with androgen substrates while HSD17B4 catalyzes the oxidative, inactivating 17β-hydroxysteroid dehydrogenase activity we observed (**Fig. 1A**), consistent with previously published findings (6). HSD11B1 expression was detectable in samples from 13/14 donors, while HSD11B2 mRNA was not detected. **Fig. 1B** schematically illustrates the enzymes identified by qPCR as in PBMCs and their roles in androgen activation and inactivation in the classic and 11-oxygenated androgen biosynthesis pathways.

**Fig. 1:**
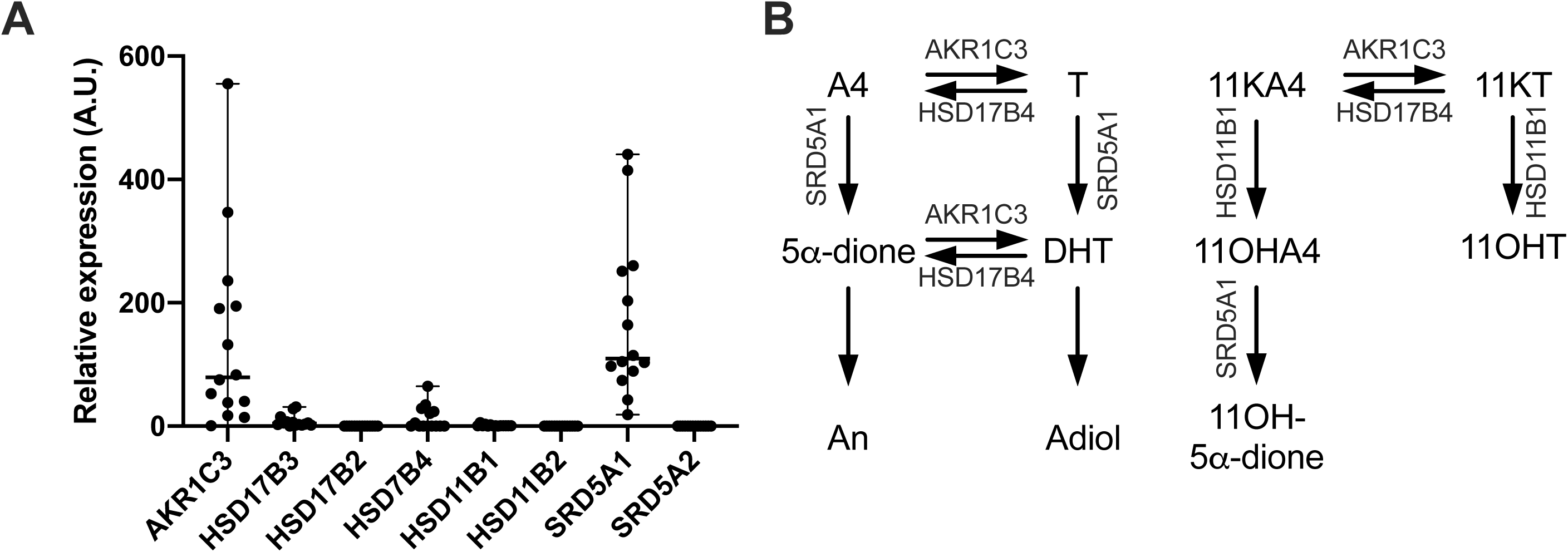
Expression of genes encoding key androgen-metabolizing enzymes in PBMCs isolated from healthy men (A, n=14; age range 22-72 years; BMI range 20.2-30.4 kg/m^2^). Gene expression was assessed by semi-quantitative PCR, normalized to HPRT1 and GAPDH and arbitrary units (A.U.) were calculates as 1000*2^-ΔCt. (B) Schematic representation of the steroid-metabolizing enzymes in PBMCs and their roles in activation and inactivation of androgens in classic and 11-oxygenated androgen pathways.

### PBMCs preferentially activate 11-oxygenated androgens

In order to unravel the pathways of androgen metabolism in PBMCs, we isolated PBMCs from male healthy donors and performed ex vivo incubations, using as substrates several androgen precursors and active androgens from both the classic (DHEA, A4 and T) and 11-oxygenated androgen pathways (11OHA4, 11OHT, 11KA4, 11KT). Product formation was quantified by LC-MS/MS.

Initial time course experiments with PBMCs isolated from three male donors (aged 22, 23 and 28 years) identified 24 hours as a suitable incubation time allowing for robust product quantification within the linear range of product formation over time (data not shown). These initial experiments additionally revealed very low 3β-hydroxysteroid dehydrogenase activity in PBMCs and thus excluded DHEA as a relevant substrate for metabolism in PBMCs (data not shown).

All further experiments were performed with PBMCs isolated from men aged 18-30 years (n=4-5; age 22-30 years; BMI 20.2-29.1 kg/m^2^) and men aged >50 years (n=4-7; age 53-72 years; BMI 21.2-30.4). Since we did not observe any significant differences in androgen metabolism and expression of genes encoding key androgen-metabolizing enzymes between the two age groups, we present and discuss the combined data for the entire cohort below (n=8-12; age 22-72; BMI 20.2-30.4 kg/m^2^).

Incubations with A4 and T yielded their respective 5α-reduced products, 5α-androstanedione (5α-dione) and 5α-dihydrotestosterone (DHT), and revealed efficient interconversion of A4 and T by reductive and oxidative 17β-hydroxysteroid dehydrogenases (**Fig. 2A+B**). 5α-dione and DHT were further converted to their 3α-hydroxy metabolites androsterone (An) and 5α-androstanediol (Adiol).

**Fig. 2:**
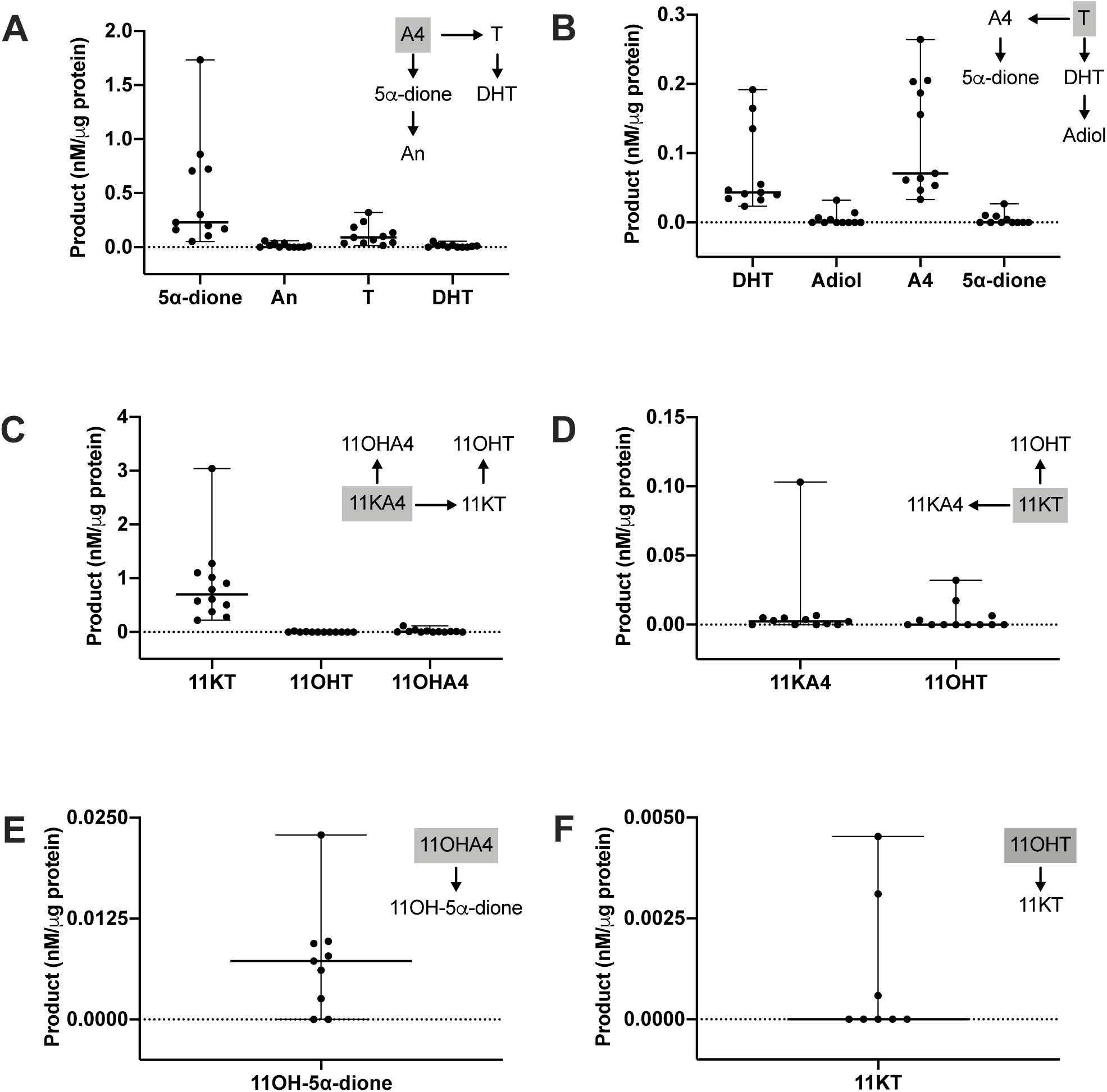
Ex vivo metabolism of classic (A+B) and 11-oxygenated androgens (C-F) by PBMCs isolated from healthy men (n=8-12; age range 22-72 years; BMI range 20.2-30.4 kg/m^2^). PBMCs were incubated with 100 nM substrate for 24 hours. The respective substrate is shown in a grey box for each graph. Product formation was quantified by LC-MS/MS and normalized to the total protein content of the incubation. Median and range are indicated. Product concentrations below the LLOQ are shown as 0.

The quantitatively dominant formation observed among all substrates tested was the generation of 11KT from 11KA4, which was significantly higher than the generation of T from A4 (p=0.001; n=11; **Fig. 2C**). After an incubation period of 24 hours, PBMCs generated approximately 8 times more 11KT than T from their respective precursors 11KA4 and A4 (**Fig. 2A+C**). However, while T was converted back to A4 in large quantities, incubation with 11KT led to only minor generation of 11KA4 (quantifiable in 8/12 incubations, **Fig. 2D**), further contributing to preferential 11KT activation by the PBMCs.

Expressing the observed steady state between activation and inactivation as product/substrate ratios made this difference even more obvious, clearly indicating that the generation of active 11-oxygenated androgens is favored in PBMCs (**Fig. 3A**). Product/substrate-ratios for different substrates of 5α-reductase reflect the established substrate preference for SRD5A1 with A4 resulting in the highest activity followed by T and with only minor activity for 11OHA4 (**Fig. 3B**).

**Fig. 3:**
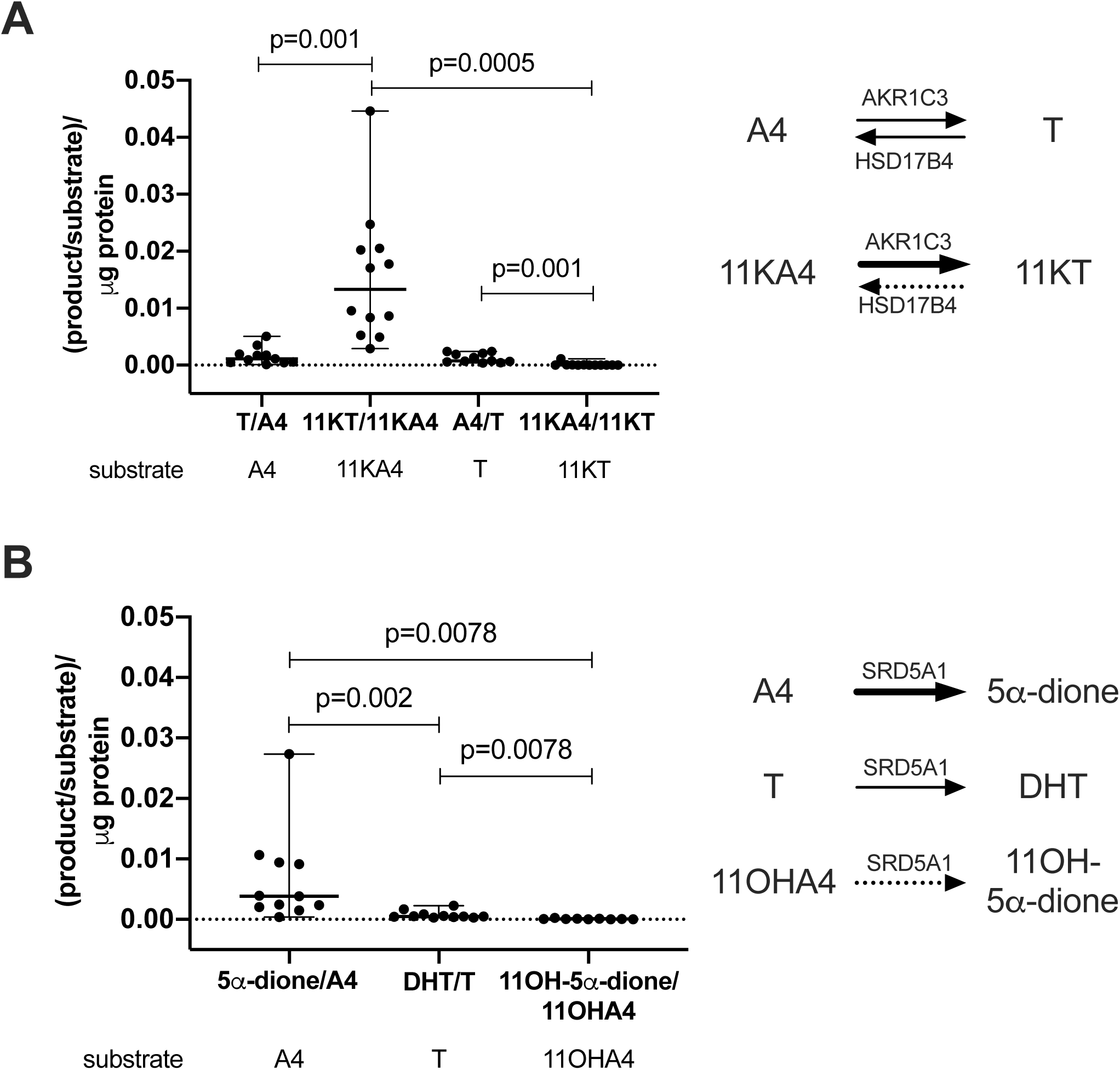
Product/substrate ratios in ex vivo incubations of PBMCs isolated from healthy men (n=8-12; age range 22-72 years; BMI range 20.2-30.4 kg/m^2^) after the addition of different steroid substrates. (A) Steady-state ratios of 17β-hydroxysteroid dehydrogenase products and substrates, reflecting the dominant contribution of the reductive, activating 17β-hydroxysteroid dehydrogenase AKR1C3 to 11-oxygenated androgen metabolism. (B) 5α-reductase activity by SRD5A1. Product and substrate were quantified by LC-MS/MS after a 24-hour incubation. Ratios were normalized to the total protein content of the incubation. Statistical analysis was performed using Wilcoxon matched-pairs signed rank test.

The generation of 11OHA4 from 11KA4 (quantifiable in 8/12 incubations) and of 11OHT from 11KT (quantifiable in 4/12 incubations) indicated 11β-hydroxysteroid dehydrogenase type 1 (HSD11B1) activity in PBMCs, however, at negligible levels compared to the 17β-hydroxysteroid dehydrogenase and 5α-reductase activities observed (**Fig. 2C+D**). We did not detect any 5α-reduced products of 11KA4 and 11KT (11-keto-5α-androstanedione and 11-keto-5α-dihydrotestosterone) (n=3; data not shown).

Incubation with the 11-oxygenated androgen precursors 11OHA4 and 11OHT led to only negligible product formation compared to the other substrates tested. After 24 hours the generation of the 5α-reduced product 11β-hydroxy-5α-androstanedione (11OH-5α-dione) from 11OHA4 could be quantified, as well as the generation of 11KT from 11OHT (**Fig. 2E+F**).

### AKR1C3 expression and activity in PBMCs is primarily driven by natural killer cells

In order to identify the subpopulation(s) within the PBMC compartment responsible for the AKR1C3-catalysed androgen activation observed in our PBMC incubations, we used publicly available RNAseq-based gene expression data from 15 FACS-sorted PBMCs subpopulations including B-cells, different T-cell populations, monocytes and natural killer (NK) cells (dice-database.org (25)). This revealed significantly higher *AKR1C3* expression in NK cells (p<0.0001) than in any other PBMC subpopulation (**Fig. 4A**). There was no difference between *AKR1C3* expression in NK cells from male and female donors (**Fig. 4B**). To confirm NK cells as the major site of AKR1C3 activity, we isolated PBMCs from a leukocyte cone of an anonymous donor and subsequently enriched NK cells in a fraction of the PBMCs. Ex vivo incubations of the matched crude PBMC isolations and the enriched NK cell fraction with the AKR1C3 substrates A4 and 11KA4 confirmed higher AKR1C3 activity in the NK cell enriched incubation compared to the incubation with crude PBMC isolates (**Fig. 4C**).

**Fig. 4:**
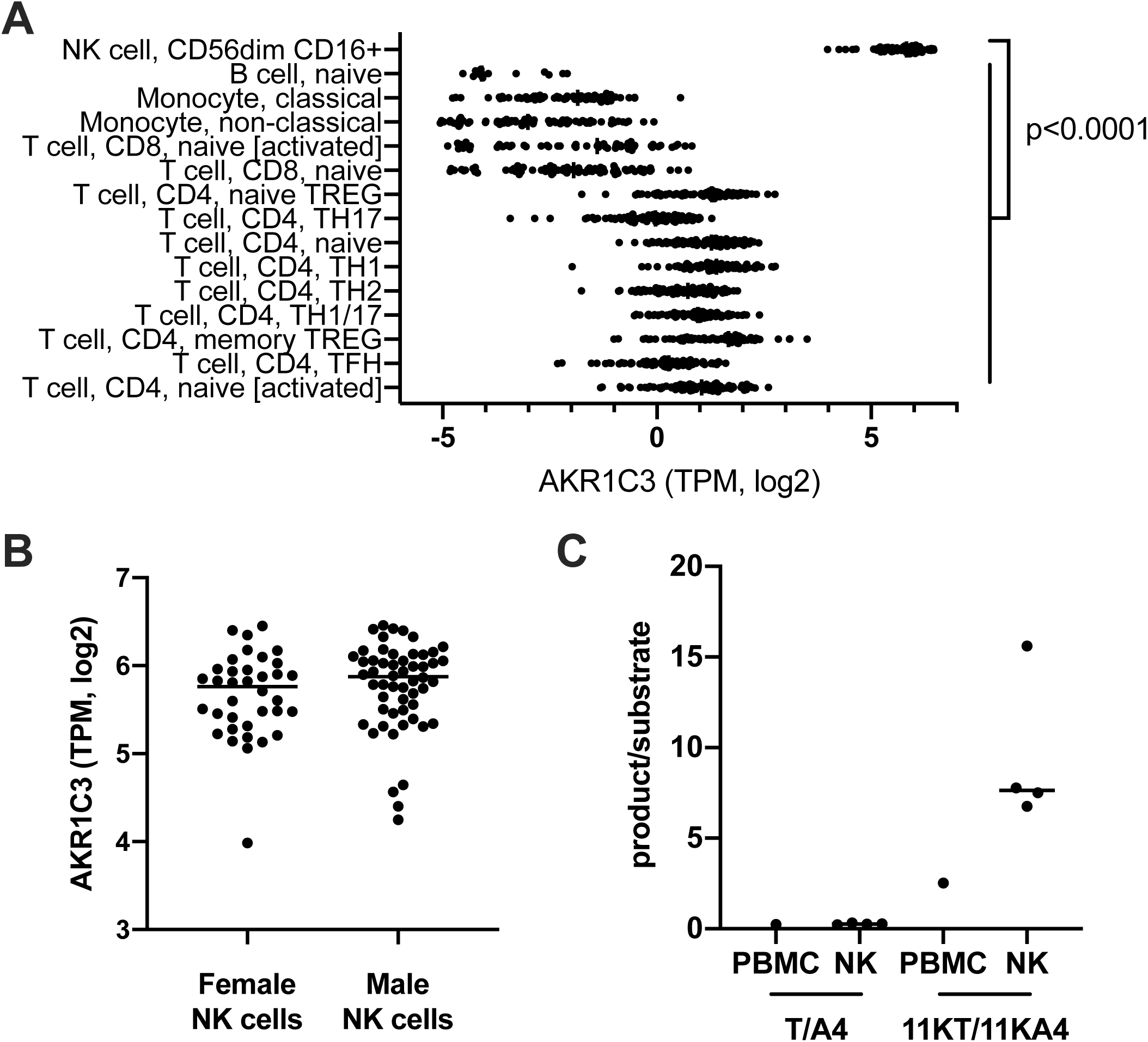
Natural killer cells are the major peripheral blood mononuclear cell population with *AKR1C3* expression (A and B) and activity (C). RNAseq analysis of FACS-sorted PBMC subpopulations (dice-database.org; (25)) revealed natural killer cells (NK cells) as the cell type with the highest *AKR1C3* expression (A) with comparable expression in women (n=37) and men (n=54) (B). NK cell-enriched incubations with AKR1C3 substrates (n=4 technical replicates) showed higher AKR1C3 activity (shown as product/substrate ratio) compared to incubations of crude PBMC preparations from the same donor (n=1) (C). NK cells enriched using MACS negative selection (Miltenyi Biotec) were incubatedand product formation after 24 hours analyzed by LC-MS/MS. Statistical analysis of *AKR1C3* expression in the PBMC subpopulation was performed by one-way ANOVA followed by Tukey’s multiple comparisons test. *AKR1C3* expression in male and female NK cells was compared by Mann-Whitney test. TPM, transcripts per million.

### Lag time prior to the separation of cellular components increases 11-ketotestosterone

We collected blood samples from six healthy volunteers (3m, 3 f; age range 28-50 years) to assess the effect of an extended incubation of their blood samples unseparated from cellular components on the quantification of serum steroids by LC-MS/MS (**Fig. 5**). We observed a time-dependent increase in the serum concentrations of the AKR1C3 product 11KT, reaching a median relative increase of 44% after 24 hours, with significant increases in 11KT observed after 2 hours of leaving the full blood samples unseparated (**Fig. 5I**). Additionally, we observed decreases in the concentrations of the AKR1C3 substrates A4 and 11KA4, with median relative decreases of 19% and 34%, respectively, at 24 hours. The observed decreases in serum A4 concentrations were statistically significant for the majority of time points assessed (p<0.05), while statistical significance was not reached for the decrease in 11KA4 (**Fig. 5C+H**).

**Fig. 5:**
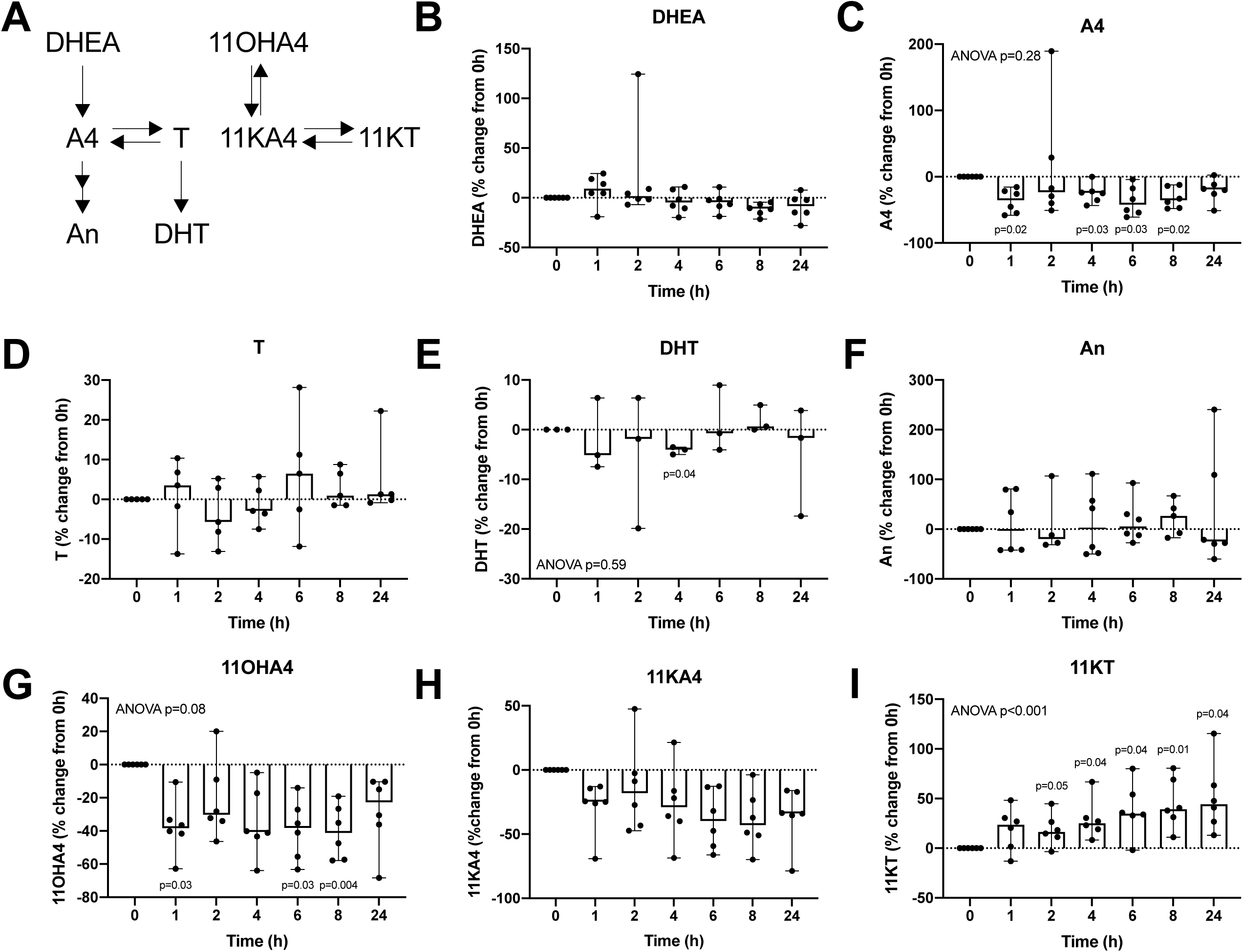
Relative changes in androgen concentrations compared to baseline, measured in serum after incubation of unseparated whole blood samples. Baseline samples were separated by centrifugation within 1h after collection. The remaining samples were incubated as whole blood at room temperature for the times indicated prior to separation of serum and cellular content (sample numbers: n=6 for DHEA (**B**), n=6 for A4 (**C**), n=5 for T (**D**), n=3 for DHT (**E**), n=6 for An (**F**), n=6 for 11OHA4 (**G**), 11KA4 (**H**) and 11KT (**I**)). Individual data points are shown with range and median. Statistical analysis was performed on the concentrations in nM using ANOVA, followed by Dunnett multiple comparison test to compare each time point against baseline.

Among the other analytes investigated (DHEA, T, DHT, An, 11OHA4), only 11OHA4 exhibited significant changes over time (p<0.05), with a median relative decrease of 38% occurring within one hour after blood collection (**Fig. 5G**).

## Discussion

Androgen signaling is vital for immune cell function by regulating proliferation, differentiation, cytokine production and other pathways (7, 8). Intracrine androgen activation from adrenal androgen precursors contributes a large proportion of androgen receptor activation in peripheral target cells and tissues of androgen action (17, 26). While we have previously shown that PBMCs can activate the classic androgen precursor A4 to T via AKR1C3 and further convert T to the most potent human androgen DHT (6), we show for the first time in this study that PBMCs preferentially activate the 11-oxygenated androgen precursor 11KA4 to its active counterpart 11KT. We show that PBMCs generate approximately 8-fold more 11KT than T from their respective precursors, revealing 11KT as the predominant active androgen within the human PBMC compartment. Using in vitro systems overexpressing AKR1C3, Barnard et al. (10) previously demonstrated that AKR1C3 has a significantly higher catalytic efficiency for the activation of 11-oxygenated androgens compared to classic androgens, which we have shown here, for the first time, ex vivo in human cells.

In addition, we show that the inactivating conversion of the active androgens T and 11KT to their respective precursors A4 and 11KA4 by oxidative 17β-hydroxysteroid dehydrogenase activity is relevant only for T, while inactivation of 11KT to 11KA4 occurs only in negligible amounts. Taken together, this demonstrates that PBMCs preferentially activate 11-oxygenated androgens. This is in agreement with the study by Barnard et al. (10) who showed that while 17β-hydroxysteroid dehydrogenase type 2 (HSD17B2) catalyzes the inactivation of T and 11KT with similar efficiencies, increased ratios of AKR1C3 to HSD17B2 favor the activation of 11-oxygenated androgens due to the catalytic preference of AKR1C3 for 11KA4 over A4.

Our results confirm previous findings that steroid 5α-reduction contributes to the activation of classic androgens in PBMCs by generating the most potent human androgen DHT from T (6). The observed substrate preference of 5α-reduction for A4 over T is consistent with the established substrate preference of steroid 5α-reductase type 1 (SRD5A1) (27), which we confirm is the major steroid 5α-reductase in PBMCs (6). We did not observe relevant 5α-reduction of 11-oxygenated androgens. Using in vitro promoter reporter assays 11KDHT has been shown to activate the AR with potency and efficacy comparable to DHT. However, it is not clear if the generation of 11KDHT by the 5α-reduction of 11KT is relevant under physiological conditions. Recently we showed that 11KT metabolism primarily proceeds via the AKR1D1 mediated 5β-reduction of the steroid A-ring and that SRD5A2, but not SRD5A1, can efficiently catalyze the 5α-reduction of 11KT (28). We now confirm that 11KT is not 5α-reduced by human cells with SRD5A1 expressed at levels that efficiently convert T to DHT, confirming that the 5α-reduction of 11KT would require the expression of SRD5A2.

In this study, we did not observe any significant effect of age on androgen activation in PBMCs with only a trend for increased median AKR1C3 and SRD5A1 activity in contrast to a published study by our group that described significantly increased AKR1C3 and SRD5A1 activity in men aged over 50 compared to men aged 18-30 years (6). The small sample numbers of both studies and differences in the assays used for steroid quantification (LC-MS/MS in our assay vs. thin layer chromatography) are likely to responsible for this discrepancy. However, when assessing the effect of age on androgen activation in peripheral tissues, age-related changes in the supply of androgen precursors from circulation need to be considered, in addition to age-dependent changes in the expression of androgen-activating and - inactivating enzymes in the peripheral target tissues of androgen action. While circulating levels of classic androgens significantly decline with age, levels of 11-oxygenated androgens remain constant across adulthood (29, 30). Hence, the peripheral activation of 11-oxygenated androgens is favored over the activation of classic androgens not only by the substrate preference of the key androgen activating enzyme AKR1C3, but also by constant high substrate availability across the life span.

Primary adrenal insufficiency is associated with an increased risk of infections compared to the general population (31). In addition, patients with primary adrenal insufficiency show significantly reduced NK cell cytotoxicity (32). Interestingly, this finding not affected by DHEA replacement therapy excluding the deficiency of the classic androgen pathway precursor DHEA as a cause of the reduced NK cell cytotoxicity. Our study identifies adrenal 11-oxygenated androgen precursors rather than classic androgen precursors as predominant androgens activated in the PBMC compartment and particularly in NK cells, potentially linking the lack of adrenal 11-oxygenated androgen precursors in primary adrenal insufficiency to the observed decrease in NK cell cytotoxicity.

The preference of PBMCs to generate 11KT from 11KA4 via AKR1C3 activity is further reflected in the significant increases in 11KT serum concentrations, if cellular components were not removed from the full blood samples in a timely fashion, confirming previous preliminary observations (23). We found that these changes became significant after two hours on benchtop without separation, suggesting that blood samples for the measurement of 11-oxygenated androgens should be processed within two hours of collection. 11KT is the dominant circulating active androgen in polycystic ovary syndrome (15) and CAH (14, 33) and a useful marker for the diagnosis of androgen excess. The ongoing activation of 11KT in unseparated full blood samples suggests that saliva, a cell-free biofluid, could be a superior matrix for the measurement of 11KT (33, 34).

In conclusion, we show that 11-oxygenated androgen precursors are the preferred substrates for androgen activation in PBMCs, yielding 11KT as the major active androgen in the PBMC compartment. This is catalyzed by AKR1C3, which is predominantly expressed in NK cells, potentially linking adrenal 11-oxgenated androgen deficiency to the reduced NK cell cytotoxicity in primary adrenal insufficiency. Androgen metabolism by PBMCs can affect the measurement results of 11-ketotestosterone serum concentrations, if the cellular component of whole blood samples is not removed in a timely fashion.

## Acknowledgments

We thank all volunteers for the donation of blood samples. We would like to thank Professors Martin Hewison and Karim Raza, University of Birmingham, for their help with obtaining ethical approval for blood cone collections. We are grateful to Emily Powell, University of Birmingham, for her help with the PBMC and NK cell isolation.

